# Regulated apoptosis is a conserved mechanism pausing female reproduction and establishes the sterile worker caste in the eusocial wasp, *Polistes*

**DOI:** 10.64898/2026.07.07.732837

**Authors:** Laura E. Miller, Ella S. McVerry, Sean O’Donnell, Kari F. Lenhart

## Abstract

Female reproduction is an energetically expensive process, so species evolve to balance survival with reproductive output. Many female organisms can temporarily pause their reproduction, including egg development, in response to physiological stress. The cellular mechanisms initiating and maintaining a stress-induced pause in oogenesis have been most extensively studied in *Drosophila melanogaster.* While the molecular control of paused oogenesis in response to starvation have been well characterized in flies, it remains unknown if these mechanisms are shared by other species with regulated pauses in oogenesis. Eusocial insects are characterized by a reproductive division of labor, with colonies of reproductive queens and sterile female workers. The social paper wasp, *Polistes*, has a dynamic dominance-based hierarchy for queen status. Worker *Polistes* are kept sterile by a combination of social and nutritional stressors. Here, we establish *Polistes* as a model to explore adult female reproductive plasticity. Through immunohistochemistry we have directly compared the *Drosophila* and *Polistes* ovarian structure and identified critical regions of the ovary in wasps that undergo regulated cell elimination during reproductive pause in flies. By comparing tissue structure, cell organization and rates of cell death between *Polistes* queens and workers we identified apoptosis as a key regulator maintaining worker sterility. Critically, this mechanism appears to be partially conserved with that in *Drosophila*. Finally, we find that changes in the timing and location of cell death in *Polistes* workers implicate oocyte identity and oocyte growth as additional potential regulators of temporary disruption of oogenesis.

**Summary Statement:** Establishing the social paper wasp, *Polistes*, as a new model for female adult reproductive plasticity via temporary pausing of oogenesis in the sterile female workers.

## Introduction

Female reproduction is costly, as egg production is much more energetically expensive than sperm production (Edward & Chapman, 2011). Therefore, selection often balances maximizing reproductive output with female longevity (Edward & Chapman, 2011). Many females from insects to mammals have mechanisms to temporarily pause reproductive ability, including egg production, in response to physiological stress, such as starvation and high temperatures (Meiselman, Kingan & Adams, 2018) (DeRensis et al., 2021) (Wang et al., 2017). Eusocial insects in the order Hymenoptera (ants, bees and wasps) are characterized by the presence of reproductive division of labor maintained by differences in female fertility (Cronin et al., 2013) (Cardoso-Júnior et al., 2026) (Tibbets et al., 2011). The colonies of eusocial insects consist of a single or multiple reproductive queens and many sterile female workers with undeveloped ovaries (Cronin et al., 2013) (Cardoso-Júnior et al., 2026) (Tibbets et al., 2011). Importantly, death of a queen induces reproductive development of one or more previously sterile female workers. Thus, eusocial Hymenoptera exhibit extreme adult reproductive plasticity and are excellent systems for analyzing mechanisms for temporary reproductive diapause at the cellular level.

Our understanding of the cellular mechanisms that transiently halt oogenesis, has been informed by studies in the fruit fly *Drosophila melanogaster* (Meiselman, Kingan & Adams, 2018) (Drummond-Barbosa & Spradling, 2001). The fly ovary consists of 16 individual ovarioles, each of which contains a germarium, housing the germline stem cells, followed by strings of progressively differentiating egg chambers (King, 1970) (Spradling, 1993) (Bastock & Johnson, 2008) (Fig.1C,D). Germline stem cells divide to produce differentiating daughter cells termed cystoblasts. These cells undergo four rounds of mitotic division with incomplete cytokinesis to generate interconnected germline cysts containing 16 cells. Fifteen of these are nurse cells that will produce RNAs and proteins to later deposit into the single oocyte (Bastock & Johnson, 2008) (Huynh & St. Johnson, 2004) (Guild et al., 1997) (Spradling, 1993). Each cyst then becomes encased by a layer of somatic follicle cells to generate an egg chamber (Fig.1A,C,D). Egg chambers continue to develop through 14 stages, ultimately forming a single oocyte.

**Figure 1.**
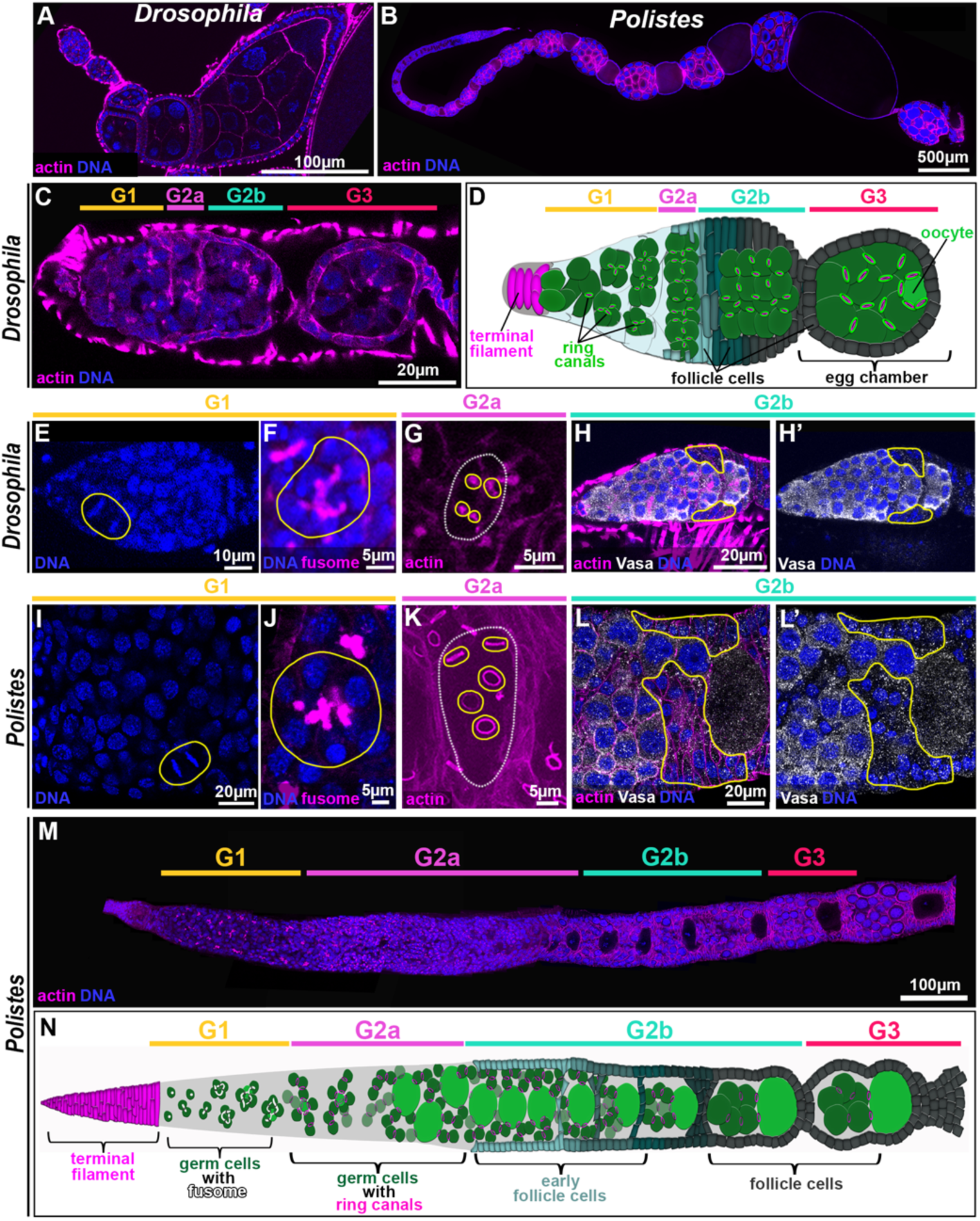
Conservation in regional identity within germaria of *Drosophila* and *Polistes*. Immunofluorescent images of *Drosophila* (A,C,E-H’) and *Polistes* (B,I-M) ovarioles with DNA stained by Hoechst in blue, actin stained by phalloidin in magenta the fusome in *Drosophila* stained by 1B1 in magenta (F) and germ cell marker Vasa in white (H-H’, L-L’). (A-B) DNA and actin staining of individual ovarioles from *Drosophila* (A) and *Polistes* queen (B). (C-D) Immunofluorescent image (C) and diagram (D) of the *Drosophila* germarium indicating regional identity from G1-G3. (E and I) Hoechst staining highlighting mitotic germ cells (yellow circle) within G1 of the fly (E) and wasp (I) germaria. (F and J) DNA and fusome staining indicating branched fusome structure extending between mitotically dividing germ cells within G1 cysts in fly (F) and wasp (J). (G and K) Actin accumulation at ring canals (yellow circles) connecting the developing oocyte (white circle) to adjacent nurse cells after completion of mitotic divisions in G2a in fly (G) and wasp (K). (H,H’ and L,L’) Vasa-positive germ cells begin to associate with Vasa-negative somatic follicle cells (yellow outline) within the G2b region of the germarium in fly (H,H’) and wasp (L,L’). (M,N) Immunofluorescent image (M) and associated diagram (N) of regional identity within the *Polistes* germarium.

Nutritional stress induces decreased reproductive output in *Drosophila* at two distinct points in oogenesis. First, starvation promotes significant apoptosis within the germarium. While all germ cells in the germarium decrease mitotic rates during starvation, apoptosis is induced very specifically in cysts at the G2a/G2b region where mitotic divisions have completed and somatic follicle cells are beginning to encase the germline cysts (Drummond-Barbosa & Spradling, 2001). Second, a critical nutritional checkpoint occurs as cysts enter stage 7/8 of oogenesis, the point at which yolk uptake (vitellogenesis) begins. While earlier developing egg chambers can survive mild starvation, lack of sufficient environmental nutrients forces degeneration at entry to stage 8 prior to the energetically costly process of vitellogenesis being initiated (Jouandin, Ghiglione, & Noselli, 2014) (Meiselman, Kingan & Adams, 2018) (Drummond-Barbosa & Spradling, 2001). Whether these cellular mechanisms regulating paused reproduction in female flies are broadly conserved features of reproductive plasticity across insects remains entirely unknown.

We have established the paper wasp *Polistes* as a model for extreme reproductive plasticity to interrogate the degree of conservation in control of paused oogenesis. *Polistes* exhibits a caste system where adult reproductive suppression is an obligate feature of reproductive division of labor. Importantly, *Polistes* have a dynamic caste system with workers that can replace the queen (O’Donnell, 1998) (Jandt & Toth, 2015). This suggests that the cellular mechanisms enforcing sterility in workers can be reversed to permit successful oogenesis just as in the temporary reproductive pause in *Drosophila* (Torres et al., 2014) (Ratnieks, 1988) (Taylor et al., 2020).

The *Polistes* caste system is maintained by a dominance hierarchy, where the most dominant individual is often the queen (Pardi, 1948) (Jandt, Tibbetts & Toth, 2014). If the queen is lost, a series of “fights” occur among workers in which larger individuals with higher juvenile hormone levels are more likely to initiate and win (Tibbetts & Izzo, 2009). Eventually, the successive winner, who also tends to have the most developed ovaries, becomes the new queen (Tibbetts & Izzo, 2009). In addition, studies suggest that *Polistes* workers are kept sterile in part by limiting access to quality food items, especially insect prey (O’Donnell et al., 2018) (O’Donnell, 1995). Interestingly, exposure of female fruit flies to juvenile hormone under starvation conditions represses the pause in oocyte development typically observed with nutrient deprivation (Terashima et al 2005). Thus, the stimuli triggering reproductive pause and reinitiation in *Drosophila* appears to be conserved in *Polistes.* (Meiselman, Kingan & Adams, 2018). We therefore asked whether sterility in *Polistes* workers is initiated and maintained by the same cellular mechanisms that pause oogenesis in starved fruit flies.

We have conducted comparative analyses between the morphological and structural organization of ovarioles in *Polistes* queens and *Drosophila* females. By leveraging the significant existing knowledge of fruit fly oogenesis, we identified analogous structural features in the wasp ovary, permitting us to identify discrete stages of oocyte development from the germarium to late-stage egg chamber formation. Through these analyses we established the specific regions along the ovariole where critical aspects of oogenesis, including mitotic divisions, somatic cell association with cysts and oocyte specification, occur. Using this characterization, we then directly interrogated whether apoptotic cell death within the germarium or later in oogenesis is a conserved feature of reproductive repression between flies and wasps. Intriguingly, we find that *Polistes* workers exhibit significantly fewer egg chambers within the germarium compared with controls. Critically, we find this decrease in egg chambers within worker ovarioles is mediated by significant apoptotic cell death induced in the G2a region of the germarium. Thus, we find that stage-specific induction of apoptosis within early egg chambers is a conserved feature of reproductive repression, indicating that caste-based reproductive plasticity shares both global regulatory and specific cellular features with stress-induced, transient cessations in oocyte development.

## Methods

### Samples

*P. exclamans* queens and workers were collected throughout the Summer of 2024 at parks and conservatories in and around Philadelphia (John Heinze Wildlife Refuge (39.892, -75.257) and Rushton Woods Preserve (39.984, -75.487). The queens were collected along with their nests as either single foundresses in early June, or along with their workers in July and August. All wasps were dissected within 24 hours of being collected. Queens were identified as having ovarioles containing mature oocytes.

*Drosophila* stocks were maintained on Bloomington Drosophila Stock Center (BDSC) standard cornmeal medium in vials or bottles. All stocks were raised at 25°C and 3–4-day old adult females were selected for experimentation.

### Immunostaining

Immunostaining was performed as previously described (Roach et al., 2025). Briefly, ovaries were dissected in Ringers solution and fixed for 30 minutes in 50% Heptane: 4% paraformaldehyde diluted in Buffer B (75mM KCl; 25mM NaCl; 3.3mM MgCl2; 16.7 mM KPO4) followed by several washes in PBSTx (1× PBS, 0.1% Triton-X 100) and blocking in 2% normal donkey serum. Ovaries were incubated in primary antibodies at 4°C at least overnight, washed several times, and then incubated in appropriate secondary antibodies for 1-hour at room temperature. After additional washes, ovaries were equilibrated in a solution of 50% glycerol and then mounted on slides in a solution of 80% glycerol. Primary antibodies: mouse anti-Orb (DSHB; Wasps-1:20, Flies-1:30) and rabbit anti-Vasa (Boster Bio; Wasps-1:1000, Flies-1:5000) rabbit anti-dcp1 (Asp215, Cell Signaling Technology; 1:1,000). Secondary antibodies: anti-rabbit 488 and anti-mouse Cy5 (Jackson ImmunoResearch; Wasps-1:200, Flies-1:300). DNA was stained using Hoechst 33342 (Wasps-1:2000, Flies-1:3000) and actin was visualized using rhodamine phalloidin (Wasps-1:200, Flies-1:300).

### Image acquisition and analysis

Immunofluorescent images were acquired using a Leica Stellaris 5 DMi8 inverted stand with tandem scanner; four power HyD spectral detectors; and HC PL APO 63×/1.4NA CS2 oil objective using LAS X software or an Olympus iX83 inverted spinning disk confocal with Hamamatsu EM-CCD camera and 60X 1.4NA silicone oil immersion objective. Images were processed in Fiji and Imaris software was used to generate 3D reconstructions of ring canal and actin tunnel structures. Figures were generated in Adobe Photoshop.

### Ring Canal Measurements

In *Polistes,* oocytes were identified as germ cells with five associated ring canals. The oocyte ring canals were measured and categorized in the following germarium regions: early G2A, late G2A, and G2b & G3. As G3 contained only one cyst, we combined G2b and G3 into one category. We combined the measurements of ring canal diameter from egg chambers outside of the germarium into a single category. The diameter of each ring canal was measured in Fiji at the widest region of the structure. All statistical tests were carried out using the R programming software.

### Quantifying Germ Cell Cysts Between Castes

In regions G2a and G2b of the germarium, we counted the number of oocytes to indicate the number of germline cysts within those regions in both *P. exclamans* queens and workers. We identified oocytes both due to their significantly larger size compared with nurse cells and, prior to the increase in oocyte size, as the only germ cells within a cyst to have five associated ring canals. All statistical tests were carried out using the R programming software.

### Quantifying Apoptosis Between Castes

Whole nests of *P. exclamans* were collected between June and September in 2024 and 2025 at John Heinze Wildlife refuge. Nests contained a single queen and several workers. Queens were again identified as those with ovarioles containing mature eggs ready to be laid. Workers contained undeveloped ovarioles lacking mature eggs. All wasps were dissected within 48 hours and were fixed according to the immunostaining protocol. Apoptotic cells were identified using an antibody against *Drosophila* Dcp1 and the specificity of Dcp1 staining of apoptotic cells was confirmed in wasps by TUNEL assay (Sigma). Germ cells were identified as those with actin ring canals within the germarium. We quantified the number of Dcp1 positive, apoptotic germ cells within each subregion of the germarium in workers and queens. All statistical tests were carried out using R software version 4.6.0 (R Core Team, 2026). All graphs were created in R using the ggplot2 function (Wickham, 2016).

## Results

### Morphology and organization of ovarioles in *Polistes* mimic that of *Drosophila*

Oogenesis in *Drosophila* occurs in a progression from the germarium which houses the germline stem cells, mitotically dividing germ cells and early cysts through multiple stages of egg chamber development, culminating in formation of a mature oocyte at stage 14 (Fig.1A,C,D; Bastock & Johnston, 2008). To determine if *Polistes* share common mechanisms of reproductive repression with *Drosophila*, we first needed to characterize the structure of wasp ovarioles and identify specific regions within the germarium and stages of egg chamber development analogous to those in fruit fly. This was particularly critical given the dramatic size difference between ovarioles in *Polistes* and *Drosophila* (Fig.1A-B).

As nutritional deprivation induces the earliest repression of oocyte development within the fruit fly germarium, we focused our morphological analyses on the early region of the wasp ovariole and used defined characteristics of fly germarium regions to assign regional identity within the wasp. In both *Drosophila* and *Polistes,* a collection of somatic cells with stacked epithelial structure resides at the apex of the ovariole (Fig.1C,D,M,N) indicating conserved presence of terminal filament between species. Similarly, in both insects a population of mitotically dividing germ cells are present adjacent to the somatic terminal filament, indicating the germline stem cell pool (Spradling, 1993; Bastock & Johnston, 2008). The germarium in *Drosophila* is partitioned into four subregions based on the division and organization of germline cysts and somatic cells (King, 1970; Fig.1C,D). Region G1 in *Drosophila* is defined by the presence of mitotically dividing germ cells as they undergo four rounds of transit-amplifying divisions with incomplete cytokinesis to generate a 16-cell cyst containing one prospective oocyte and fifteen nurse cells (Bastock & Johnston, 2008; Fig.1C-E). During these mitotic divisions in flies, formation of stable ring canals at intercellular bridges occurs along with extension of an ER-like membranous organelle termed the fusome through each ring canal (Fig.1C-F). We determined the region within the *Polistes* germarium corresponding to G1 by the combined presence of mitotically dividing germ cells (Fig.1I, yellow outline) and presence of an extended fusome between intercellular bridges (Fig.1J yellow outline). In *Drosophila,* completion of transit-amplifying mitotic divisions within a cyst denotes the beginning of region G2a. This is visualized by the presence of a prospective oocyte with four ring canals (Fig.1D,G). We find that germ cell cysts in *Polistes* undergo an additional round of mitotic divisions within G1, leading to formation of a 32-cell cyst and a prospective oocyte with five associated ring canals (Fig.1K). The G2a/G2b transition in *Drosophila* is characterized by the initial association of somatic follicle cells with the early meiotic germline cyst (Fig.1D, yellow outline H,H’). By staining for the germ cell marker Vasa, we identified the region in the *Polistes* germarium in which mitotic germ cell divisions have completed, and Vasa-positive germ cells begin to associate with Vasa-negative cells with smaller nuclei, indicative of somatic follicle cells (Fig.1L,L’, yellow outline).

Within the G3 region in the fly germarium, the oocyte is positioned toward the posterior end of the cyst and follicle cells begin to physically separate the germline cyst from the germarium to form the first egg chamber (Figure 1C,D). We identified nearly identical changes in oocyte positioning and organization of somatic cells within regions of the *Polistes* germarium, permitting us to assign regional identity in the wasp ovariole from G1 through G3 (Fig.1M,N).

While the overall organization of the germarium and progression between differentiation stages is similar between *Drosophila* and *Polistes*, we did note several features unique to the wasp ovary. In fruit flies, oocytes and nurse cells remain similar in size until midway through oogenesis, around stage 8 (Gates, 2012; Haigo & Builder, 2011; Jia et al., 2016). Only at this point does the oocyte dramatically increase in size relative to nurse cells. By contrast, we observed visually distinctive, large oocytes in *Polistes* as early as the G2a region, suggesting that oocyte growth and, potentially, maturation is accelerated in the wasp (Fig.1M). In addition, unlike *Drosophila* in which somatic cells form a follicular epithelium encapsulating the germline cyst while remaining on the exterior of the egg chamber, we observed somatic cells intermingling with nurse cells in the interior of egg chambers throughout the length of the *Polistes* ovariole (Fig.1D,H-H’, L-L’).

### Conservation and divergence in mechanisms of oocyte specification between Drosophila *and* Polistes

The dramatic increase in oocyte size early within the *Polistes* germarium made us question whether oocyte specification may also occur more rapidly within the wasp. In *Drosophila*, oocyte identity is established and maintained by the RNA binding protein Orb (Barr et al., 2019; Fig.2A). Initially, Orb is present in all 16 germ cells within the cyst (Fig.2C,C’, yellow outline) but becomes progressively localized in late G2a/early G2b to the two pro-oocytes (Barr et al., 2019). Ultimately, Orb becomes restricted to only one germ cell of the cyst and is retained in the developing oocyte (FigE, E’, blue arrowhead). Just as in *Drosophila*, Orb is initially expressed equally in all cells within a cyst in late G2a (Fig.2D,D’, yellow outline), but then rapidly localizes exclusively to the oocyte (Fig.2D, D’,F, F’, blue arrowheads). Consistent with the timing of oocyte specification in flies, Orb becomes restricted to the pro-oocyte in *Polistes* concurrent with the oocyte positioning to the center of the ovariole, but prior to follicle cells association with the cysts (Fig.2G). Critically, although there is conservation in the dynamics of Orb expression and restriction during oocyte specification between flies and wasps, the dramatic increase in oocyte size evident in G2a of the *Polistes* germarium precedes induction of Orb in the germ cells (Fig.2D,D’, yellow arrowheads). This indicates that a distinct process initiates oocyte specification in the wasp. Critically, as we do not observe a difference in the dynamics of oocyte growth and specification between queen and worker ovaries (Fig.S1), this separate mechanism of oocyte specification does not appear to be a node of control for reproductive plasticity.

**Figure 2.**
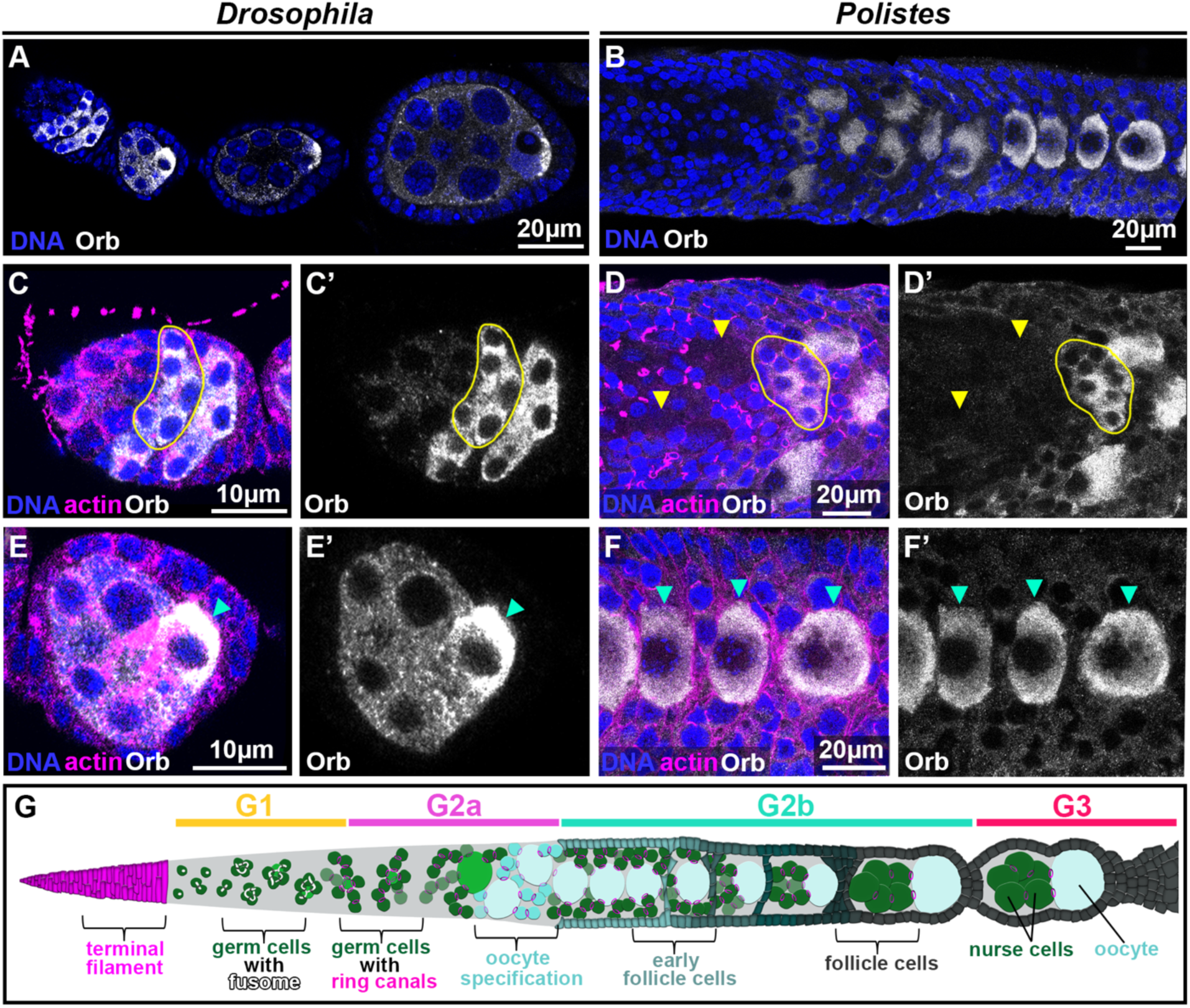
Conservation and divergence in oocyte specification between *Drosophila* and *Polistes*. (A-F’) Immunofluorescence images of actin (magenta) DNA (blue and Orb (white) in *Drosophila* (A,C-C’,E-E’) and *Polistes* (B,D-D’,F-F’). Orb is initially expressed in all germ cells of a cyst in region G2a (yellow outline, C-C’, D-D’) but rapidly becomes restricted to and retained within the prospective oocyte in region G3 (blue arrowheads D-D’ and F-F’) in both fly (A,C-C’,E-E’) and wasp (B,D-D’,F-F’). While the prospective oocyte and nurse cells are equivalent size within the germarium in *Drosophila* (C-C’, E-E’), prospective oocytes in *Polistes* are significantly larger than associated nurse cells (D-D’, F-F’), even prior to induction of Orb expression in G2a (yellow arrowheads D-D’). (G) Diagram of regional identity oocyte specification within a *Polistes* germarium.

### Interconnections between germline cysts mediating nurse cell communication with the oocyte differ between *Drosophila* and *Polistes*

In all species from *Hydra* to humans, differentiating germ cells undergo multiple rounds of mitosis with incomplete cytokinesis to establish stable intercellular bridges called ring canals. In the *Drosophila* ovary, these mitotic divisions result in formation of a 16-cell cyst with the oocyte connected to nurse cells via four stable ring canals. As cysts continue to differentiate, these actin-based ring canals grow dramatically in size from stages 2-10 to permit the eventual transport of nurse cell cytoplasmic components into the oocyte to support early embryonic development after fertilization (Bastock & Johnson, 2008; Huynh & Johnston, 2004; Ong, Foote & Tan, 2010; Tilney, Tilney & Guild, 1996; Ong & Tan, 2010; Jia et al., 2016; Jackson, Alsous, & Martin, 2023) (Fig.3A-C). This “nurse cell dumping” results in a dramatic increase in oocyte size and the concurrent reabsorption of nurse cells to yield a single, large oocyte (Jia et al., 2016) (Jackson, Alsous, & Martin, 2023).

**Figure 3.**
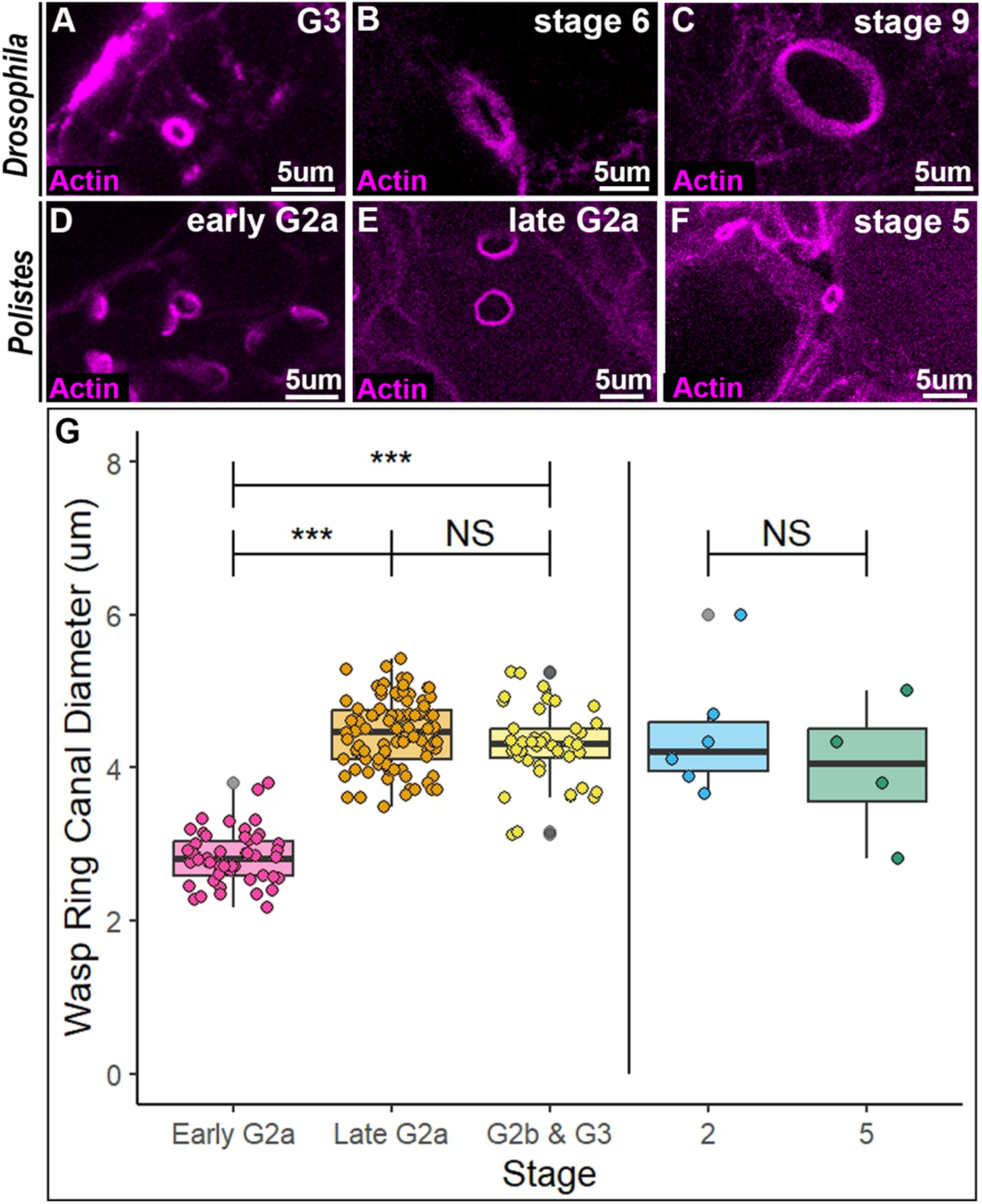
Limited expansion of ring canal diameter in *Polistes*. (A-C) Ring canals increase in diameter in the *Drosophila* ovary, with a 10-fold expansion from G3 to stage 8/9 egg chambers. By contrast, ring canal diameter in Polistes undergoes moderate expansion between early and late G2a (D,E) but ring canal size remains stable beyond G3 and overall expands far less significantly than ring canals in the fly (C,F). (G) Boxplot of ring canal diameter at different stages of oogenesis in the germarium (Early G2a, Late G2a, G2b & G3) and egg chambers (2. 5) of *Polistes* queens. N=6 ovarioles. Germarium: Anova (F value=239.1, P value= <2.2e-16) Tukey test performed between groups (***= p<0.001, NS= p≥0.05). Egg Chambers: ANOVA (F value =0.965, P value =0.4506) Tukey performed between groups.

Given the early and rapid increase in oocyte size in *Polistes*, we examined whether this process is driven by precocious expansion of ring canal diameter compared with *Drosophila*. Interestingly, while ring canals do slightly increase in size from early to late G2a in the wasp germarium (Fig.3D,E) there is no further increase in ring canal diameter during later stages of oogenesis (Fig.3F,G). Indeed, quantification of ring canal diameter between G3 and later egg chambers found no significant difference in ring canal size (Fig.2G). Importantly, however, the moderate expansion of ring canals occurs in *Polistes* immediately prior to the increase in size of the prospective oocyte, suggesting that early oocyte specification in the wasp is mediated, at least in part, by increased cytoplasmic transport from nurse cells to the oocyte.

In *Drosophila*, ring canal diameter increases 10-fold over the course of egg chamber development, with this expansion being critical for proper sharing of organelles, RNAs and proteins between the nurse cells and oocyte (Cooley, 1998). By contrast, we observed less than a 2-fold change in ring canal diameter during oogenesis in *Polistes*. This suggests that persistent and robust nurse cell-to-oocyte communication must be achieved in the wasp via a distinct mechanism.

In *Drosophila*, the oocyte is always in direct contact with the four closest nurse cells within the egg chamber maintaining connections via the largely expanded ring canals (Logan, Chou, & McCartney, 2022; Fig.4A-C). By contrast, we find that oocytes become physically separated from nurse cells in *Polistes* egg chambers, instead maintaining connection via formation of an actin-based tunnel structure (Fig.4D-I). This tunnel forms early in oogenesis, consistently visible between the oocyte and adjacent nurse cells in the first or second egg chamber outside of the germarium. Each nurse cell initially forms an individual actin-based connection with the oocyte (Fig.4D-F). Over time, these independent tunnels merge to generate a central tube connecting all adjacent nurse cells with the developing oocyte in a manner distinct from the ring canal associations established in the germarium (Fig.4G-I). Thus, wasp and fly engage distinct mechanisms to ensure robust communication and cytoplasmic sharing between nurse cells and the oocyte.

**Figure 4.**
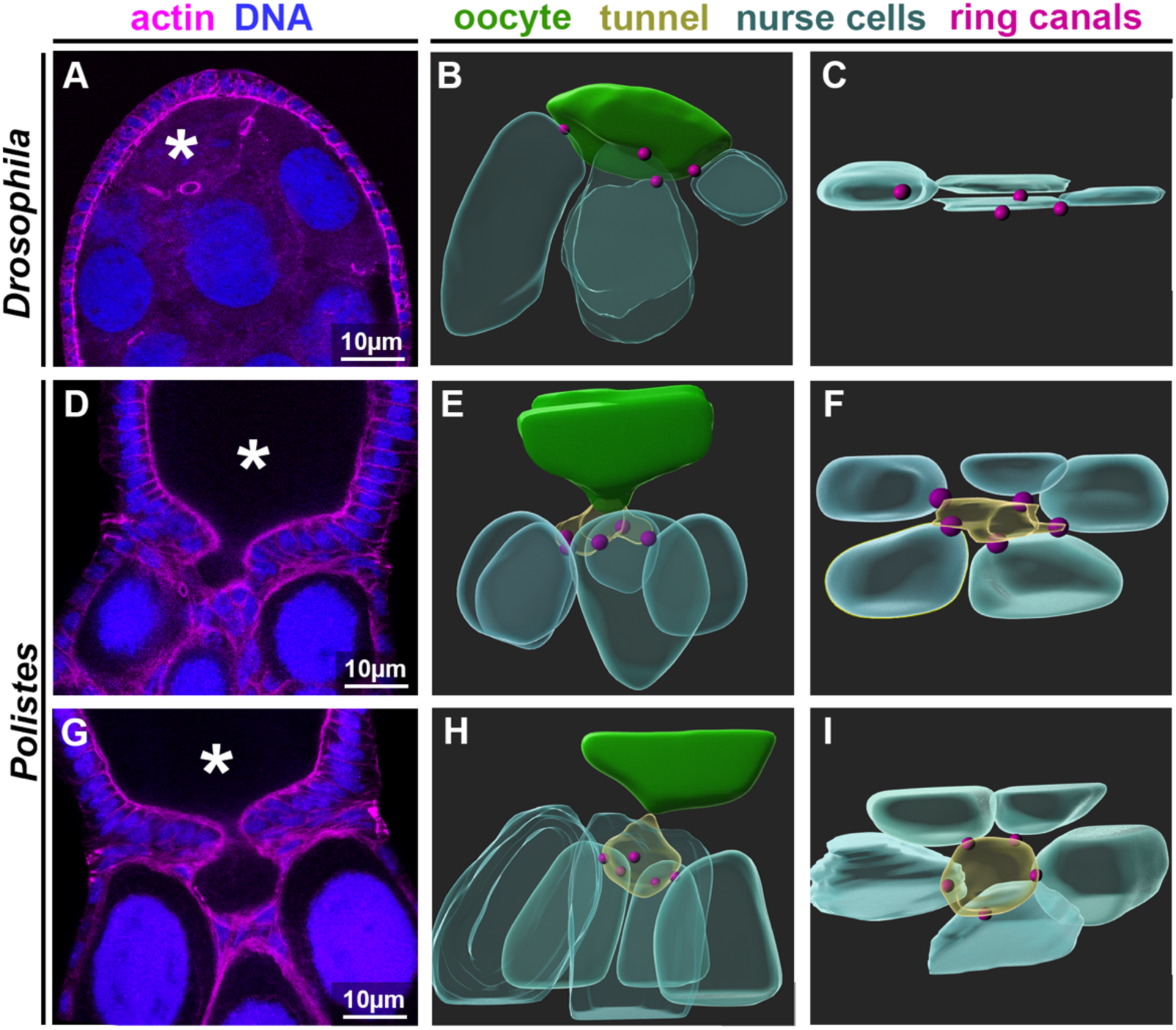
An actin tunnel connects the oocyte with nurse cells to mediate cytoplasmic sharing in *Polistes*. (A) Immunofluorescence image of a stage 6 egg chamber in *Drosophila*. (B,C,E,F,H,I) 3D reconstructions generated from immunofluorescent images of fly (B,C) and wasp (E,F,H,I) egg chambers showing nurse cells (cyan), actin ring canals (pink circles), and oocyte (green). (C) Top-down view of nurse cells connected to the *Drosophila* oocyte via ring canals (D) Immunofluorescence of initial formation of an actin tunnel connecting the oocyte with adjacent nurse cells in an egg chamber of a Polistes queen (N=4/4). (F) Top-down view of the actin tunnel and its formation relative to the existing ring canals connecting to the five closest nurse cells with the oocyte. (G) Immunofluorescence of the actin tunnel in a later egg chamber in a *Polistes* queen. (H) 3D reconstruction and modeling of the late-stage actin tunnel and (I) top-down view revealing the organization of nurse cells and ring canals relative to the later stage tunnel.

### Conservation between *Drosophila* and *Polistes* in regulation of reproductive plasticity

Having characterized the regional identity of germaria in *Polistes* queens and confirmed that general morphology and development of wasp germline cysts is similar to that in flies through the G3 stage, we next turned to comparisons between *Polistes* queens and workers to interrogate whether the mechanism of starvation-induced reproductive repression in *Drosophila* is conserved to mediate the caste-based reproductive repression in *Polistes*. There are dramatic morphological differences between ovarioles in reproductive *Polistes* queens and sterile workers (Fig.5A). Indeed, compared with the extensive string of differentiating egg chambers in the queen (Fig5A, top), only an average of one or two egg chambers are present outside of the shortened germaria region in workers (Fig.5A, bottom). We find that the overall organization of germ cell cysts within the germarium is similar between queens and workers but with workers containing significantly fewer cysts beginning in the G2a/G2b region (Fig.5B-D).

**Figure 5.**
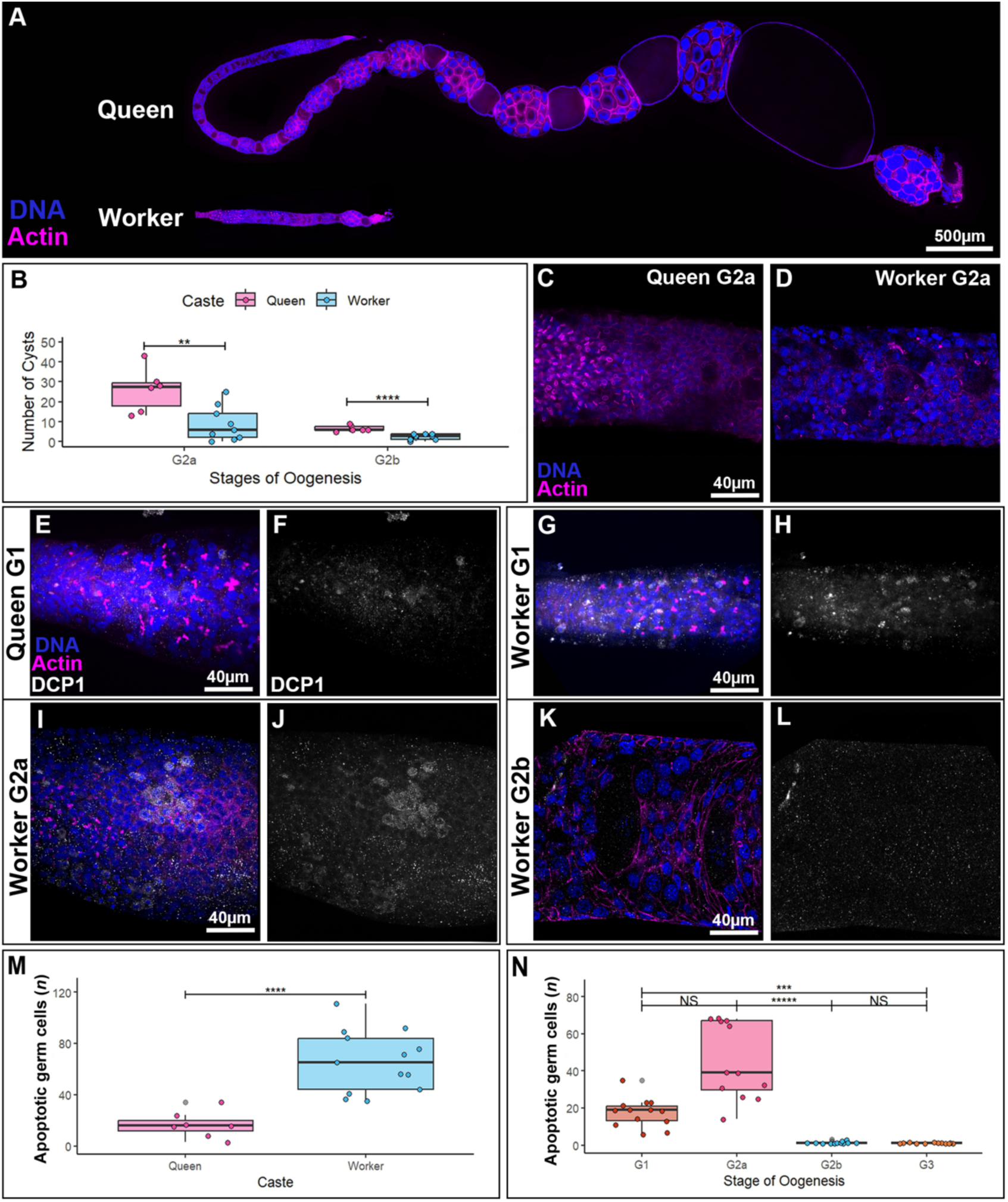
A conserved checkpoint regulates reproductive repression at the G2a/G2b transition in starved *Drosophila* and *Polistes* workers. (A) Significant morphological differences and degree of development in ovarioles from a *Polistes* queen (top) and worker (bottom). (B) Quantification of germline cyst numbers in the G2a and G2b regions of *Polistes* queen and worker germaria. G2a: N=15, T-test (t=3.3667, df=13, P-value=0.005055, **). G2b: N=15, T-test (t=5.3131, df=13, P-value=0.0001407, ****). (C,D) General morphology and presence of visibly distinct, large oocytes within germline cysts are similar between germaria of *Polistes* queens (C) and workers (D). (E-L) Dcp1 staining to detect apoptotic germ cells within the germaria of queens (E,F) and workers (G-L). G1 region of the ovaries. (M) Quantification of total numbers of apoptotic germ cells throughout the germaria of queen and worker *Polistes*. N=20, T-test (t=5.1882, df=19, p-value=6.189e-5) (N) Quantification of apoptotic germ cells within distinct regions of the germarium in workers, identifying G2a as the primary location in which cell death is induced N=12, Kruskal-wallis test (χ^2^=46.426, df=3, p-value=4.604e-10) Dunn test performed between groups: (G1-G2a: p-value=0.2981, NS, G2a-G2b: p-value=3.758e-7,*****, G1-G3, p-value=8.585e-4, ***, G2b-G3: p-value=1.000, NS)

To determine if this extreme repression of egg chamber production in workers is mediated by the same cellular mechanisms as nutrient-dependent repression in *Drosophila*, we quantified germ cell apoptosis within germaria of *Polistes* queens and workers. In starved female *Drosophila*, germline cysts undergo apoptosis at the transition between regions G2a and G2b of the germarium, with little cell death evident elsewhere within the germarium (Drummond-Barbosa & Spradling, 2001) (Pritchett et al., 2009). Interestingly, we find that *Polistes* workers have significantly more apoptotic germ cells than queens (Fig.5E-H, M), indicating that programmed cell death likely plays a role in worker sterility. To determine if germ cell death is initiated at the same checkpoint within the germarium in worker *Polistes* as in starved *Drosophila*, we quantified apoptotic germ cells at specific regions of the germarium in workers (Fig.5N). Excitingly, we find that the majority of apoptotic germ cells are located within the G2a region of the worker germarium and are not present in G2b (Figure 5E-N). This strongly indicates a conserved checkpoint for initiating and ceasing apoptotic germ cell death between *Polistes* workers and the nutritionally dependent response in *Drosophila*. However, in *Polistes* workers there is also a significant number of apoptotic germ cells within region G1, indicating that the cell death in *Polistes* is more pervasive and initiates earlier than in starved *Drosophila* (Figure G-H, N). These data strongly suggest a conserved mechanism for induction of reproductive repression between *Drosophila* and *Polistes*, with a similar checkpoint engaged during both transient (fly) and developmentally regulated (wasp) disruption of oogenesis.

## Discussion

By leveraging the extensive knowledge of *Drosophila* oogenesis, we have taken a comparative analysis approach to establish the paper wasp *Polistes* as a new model to interrogate the degree of conservation in cellular mechanisms regulating reproductive plasticity. We found that morphological and developmental features of the wasp ovariole are similar to that in fruit fly. We used discrete cellular features of oocyte development in *Drosophila* to assign regional identity to oogenesis within the germarium of *Polistes* queens. Interestingly, while Orb localization during oocyte specification and the generation of interconnected cysts via ring canals are conserved across flies and wasps, we identified several unique features of oocyte development in *Polistes*. We found that oocytes undergo significant growth early in the wasp germarium, with oocytes becoming visibly distinct from surrounding nurse cells *before* Orb induction and molecular specification of the oocyte. This precocious oocyte growth is likely mediated by an expansion of ring canal diameter that occurs between the early and late G2a regions, immediately prior to the development of visibly distinct oocytes within a germline cyst.

Additionally, we found that nurse cell-oocyte communication is likely regulated in *Polistes* via a distinct mechanism. Rather than cytoplasmic and organelle sharing being controlled through significant expansion of ring canal diameter, as in *Drosophila*, *Polistes* oocytes generate actin-based tunnels with adjacent nurse cells that coalesce into a large, singular tunnel connecting the supporting germ cells to the oocyte.

Most importantly, we leveraged our comparative analyses and characterization of *Polistes* oogenesis to address the cellular mechanisms underpinning repression of reproduction in this caste-based system. Strikingly, we found that the mechanisms to transiently pause oogenesis in starved *Drosophila* are conserved in the persistent reproductive repression of worker *Polistes*. Specifically, we found that the checkpoint for oocyte development within *Drosophila* germaria prior to the transition from G2a to G2b is precisely the point at which we observed significant cell death within germ cells in worker versus queen wasp ovaries. Taken together, this work identified both conserved and novel aspects of oocyte development in the eusocial wasp, *Polistes*, and established examination of worker versus queen germ cell biology as a new and powerful system to study mechanisms of reproductive plasticity.

### Conserved and divergent mechanisms for oocyte specification

In *Drosophila*, initial specification of the oocyte is driven by a combination of molecular and biophysical properties. While expression and restriction of the RNA-binding protein Orb is a key indicator of oocyte identity, several earlier events help to define the pro-oocyte within the germarium. Two of the 16 cells within a cyst initiate meiosis in region G2a, prior to Orb induction, through formation of the synaptonemal complex- yet this complex is only retained within a single pro-oocyte in G2b (Ables, 2015) (Von Stetina & Orr-Weaver, 2011) (Takeo et al., 2011) (King, 1970). The initial difference between the pro-oocytes and prospective nurse cells is thought to be specified via organelle and cytoskeletal asymmetries. From the first mitotic division, the prospective oocyte maintains a longer centriole than its daughter cells (Riparbelli, Persico, & Callaini, 2021) and inherits an asymmetrically greatly proportion of fusome material (Lin & Spradling, 1995) (Barr et al., 2024) (Cabrita & Martinho, 2023). As oocyte specification requires the fusome to anchor *orb* mRNA and protein in the two pro-oocytes, these early organelle asymmetries are thought to drive initiation of oocyte specification prior to Orb induction (Barr et al., 2024).

A similar combination of factors appears to be at play to specify oocyte identity in the *Polistes* germarium. Just as in flies, Orb protein is induced in all cells of the germline cysts but rapidly localizes to the prospective oocyte. And, just as in fruit flies, the pro-oocyte in wasps is initially specified prior to Orb induction. In *Polistes*, this early oocyte identity is evidenced not in cytoskeletal or organelle asymmetries but in the size of the oocyte relative to associated nurse cells. Our quantification of ring canal diameters throughout the wasp ovariole suggests that ring canal expansion occurs immediately prior to asymmetric oocyte growth, suggesting that early specification and growth of *Polistes* oocytes may be induced by increased cytoplasmic sharing from nurse cells. A number of F-actin regulators are known to control ring canal expansion in *Drosophila* including Arp2/3 and formins (Hudson & Cooley, 2002) (Thestrup et al., 2020). In the future, it will be interesting to interrogate whether similar mechanisms control ring canal growth in *Polistes* and whether disruption of ring canal expansion prevents initial oocyte specification in the wasp.

### Distinct mechanisms control nurse cell-oocyte interactions in *Polistes*

Direct contact between nurse cells and oocytes is an essential feature of reproduction in *Drosophila*. This communication is maintained throughout oogenesis via ring canals. In *Polistes*, we instead found that a distinct structure, an actin-based tunnel, forms between the nurse cells and oocyte to support interactions in mid-oogenesis egg chambers.

There are three types of oogenesis in insects: panoistic, polytrophic meroistic, and telotrophic meroistic (Büning, 2006). The oocyte being accompanied by its nurse cells throughout oogenesis as seen in both *Drosophila* and *Polistes* is a characteristic of polytrophic meroistic oogenesis (Büning, 2006). However, *Polistes* may share some characteristics with telotrophic oogenesis. In telotrophic meroistic ovaries, the oocyte receives nutritional support from nurse cells, but these nurse cells remain in the germarium of the ovariole as the oocyte descends (Büning, 2006). A nutritive chord connects the oocyte to the nurse cells and extends as the oocyte develops and travels through the ovariole (Büning, 2006) (Huebner & Gutzeit, 1986). Each oocyte has an individual chord that extends to the germarium and transfers RNA from the nurse cells to the oocyte as they progress through the ovariole (Macgregor & Stebbings, 1970). The nutritive chord is made up of microtubules and is surrounded by an F-actin meshwork which resembles the *Polistes’* tunnel we identified (Huebner & Gutzeit, 1986).

The tunnel may have evolved similarly to the nutrient chord to optimize the transfer of cytoplasmic contents from nurse cells to the oocyte. The polytrophic meroistic system relies on the constant rearrangement of microtubules (Therkauf et al., 1992) (Lu et al., 2022) and actin cytoskeleton (Riparbelli & Callaini, 1995) (Bernard, Lepesant & Guichet, 2018) to facilitate cytoplasmic transfer to the oocyte. Alternatively, the actin tunnel may be an example of convergent evolution. Identifying other species that have this trait within Hymenoptera would provide this answer and further elucidate the functions of the tunnel.

### Cellular mechanisms controlling reproductive repression are conserved

We observe significant, caste-based differences in apoptotic cell death within *Polistes* germaria. Strikingly, the majority of apoptosis occurs in workers within the G2a region-precisely the developmental timepoint at which nutrient deprivation induces an apoptotic checkpoint in *Drosophila*. This suggests that both transient (starvation) and persistent (worker caste) environmental factors impinge upon similar cellular mechanisms to promote reproductive plasticity across species. In future work, we will interrogate whether induction of reproductive development in worker wasps occurs with a concurrent loss of apoptosis within the G2a region of the germarium. In addition, it will be critical to determine the molecular mechanisms driving the G2a checkpoint and whether, as in *Drosophila*, this is controlled via insulin signaling and/or juvenile hormone production.

In addition to the substantial cell death occurring within G2a, we also observed significant differences between worker and queen germaria in rates of apoptosis within G1. Intriguingly, this region, which houses the germline stem cells and early mitotic germ cells, does not exhibit significant cell death during starvation in *Drosophila*. While these data might indicate more generally pervasive germ cell death in *Polistes* to mediate the extreme reproductive repression within workers, we observe minimal germ cell apoptosis in G2b or G3. Instead, we hypothesize that the increased cell death within G1 of the *Polistes* worker germaria may be a consequence of the early and rapid oocyte growth evident in early G2a of the *Polistes* queens. In starved *Drosophila*, the G2a/G2b apoptotic checkpoint is only engaged under extreme nutrient deprivation. By contrast, the degradation of egg chambers at the later vitellogenic nutrient checkpoint occurs even under moderate starvation conditions. This is to avoid expending the metabolic energy of vitellogenesis and oocyte growth when nutritionally stressed (Shimada et al., 2011). As the prospective oocyte grows substantially in size as early as the G2a region in *Polistes*, it is possible that a similar and more stringent cell death checkpoint may exist in the G1 region to limit the metabolic energy of producing cysts that will quickly need cytoplasmic nutrients to grow.

*Polistes* workers likely share similar mechanisms for maintaining paused oogenesis with *Drosophila*. Juvenile hormone maintains its role as a gonadotropin in *Polistes*, and its levels are positively correlated with both ovary development and status in the dominance hierarchy (Tibbetts, Levy & Donajkowski, 2011) (Tibbetts & Izzo, 2009). In colonies with active queens, workers are kept from developing their ovaries and egg laying by both dominance behaviors from the current queen and policing by their fellow workers (Reeve, 1991) (Ratnieks, 1988). These dominance interactions experienced by workers likely limits their access to adequate nutrition to develop their ovaries (O’Donnell et al., 2018). Workers forage for nectar (carbohydrate) and prey (protein and lipids) but subsequently give away protein sources and most of the nectar upon their return to the nest (O’Donnell, 1995). The nutritional aspect, the role of juvenile hormone and our confirmation of the role of apoptotic germ cell death in early oogenesis strongly indicates that the mechanisms that prompt sterility in *Drosophila* may have been co-opted to create the sterile worker caste in *Polistes*.

Understanding the cellular biology behind oogenesis and what pauses oogenesis in *Polistes* could explain how eusocial insects, when transitioning from a solitary or sub-social lifestyle, co-opted a universal ovarian stress response. Furthermore, *Polistes s*terility could present novel mechanisms to temporarily disrupt oogenesis in response to environmental stressors and seasonally regulated reproductive diapause. In essence, establishing *Polistes* as a new model for temporary disruption of oogenesis could expand the understanding of the field on how the reproductive biology of female organisms respond to different external factors to maintain the balance between conserving energy and producing offspring and will provide new insights into the mechanisms regulating reproductive plasticity across species.

## Acknowledgements

This work was supported by the Robert L. and Louise B. Jeanne Social Wasp Research Grant from the International Union for the Study of Social Insects (IUSSI) North American Section (to L.E.M.) and the National Institutes of Health (R01 GM138705 to K.F.L). The image of fusome morphology in the *Drosophila* germarium in Figure 1 was generously provided by Ph.D. candidate Ido Keren.

