## Supplemental Figure for "Regulated apoptosis is a conserved mechanism pausing female reproduction and establishes the sterile worker caste in the eusocial wasp, *Polistes*"

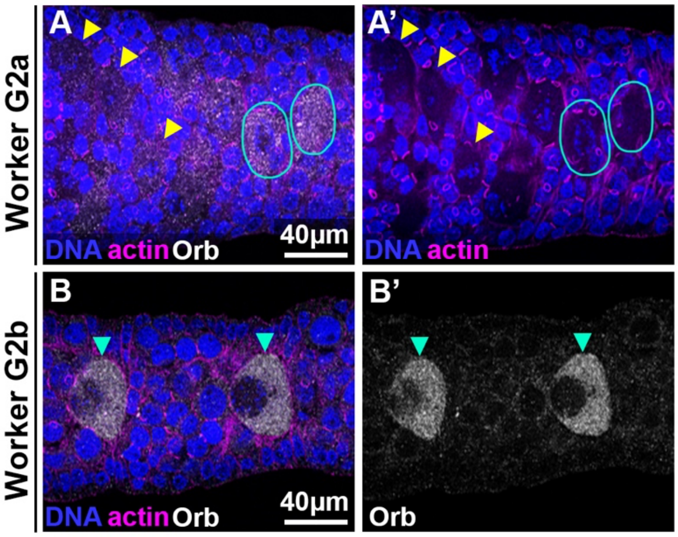


**Supplemental Figure 1. Early oocyte growth and specification remain intact in the *Polistes* worker caste.** Immunofluorescent images of DNA (Hoechst, Blue), actin (phalloidin, magenta) and Orb (gray) in the G2a (A,A') and G2b (B,B') regions of germarium. Orb is induced in the G2a region (A-A', blue circles) and retained through the G2b region (B,B', blue arrowheads) just as in ovarioles in the *Polistes* queen (main text Fig.2). In addition, workers retain the early growth and specification of oocytes within G2a prior to Orb induction (A,A', yellow arrowheads).
